# Y chromosome sequencing data suggests dual paths of haplogroup N1a1 into Finland

**DOI:** 10.1101/2024.02.23.581727

**Authors:** Annina Preussner, Jaakko Leinonen, Juha Riikonen, Matti Pirinen, Taru Tukiainen

**Affiliations:** Institute for Molecular Medicine Finland (FIMM), Helsinki, Finland; Department of Public Health, Faculty of Medicine, University of Helsinki, Helsinki, Finland; Department of Mathematics and Statistics, University of Helsinki, Helsinki, Finland

**Keywords:** Y chromosome, Y-chromosomal haplogroups, chrY, Finnish population, N1a1

## Abstract

The paternally inherited Y chromosome is highly informative of genetic ancestry, therefore making it useful in studies of population history. In Finland, two Y- chromosomal haplogroups reveal the major substructure of the population: N1a1 (TAT) enriched in the northeast and I1a (M253) in the southwest, suggested to reflect eastern and western ancestry contributions to the population. Yet, beyond these major Y-chromosomal lineages, the distribution of finer-scale Y- chromosomal variation has not been assessed in Finland. Here we provide the most comprehensive Y-chromosomal study among the Finns up to date, exploiting full sequences for 1,802 geographically mapped Finnish Y chromosomes from the FINRISK project. We assessed the distribution of common Y-chromosomal haplogroups (frequency ≥ 1%) throughout 19 Finnish regions, and further compared the autosomal genetic backgrounds of the Y-chromosomal haplogroups. With such high-resolution data, we identified novel sublineages and geographical enrichment patterns among the major Finnish haplogroups N1a1 (64%), I1a (25%), R1a (4.3%), and R1b (4.8%). Most notably, we discovered that haplogroup N1a1 splits into three major lineages within the country. While two of the sublineages followed a northeastern enrichment pattern observed for N1a1 in general, the sublineage N1a1a1a1a1a (CTS2929) (22% of all samples) displayed an enrichment in the southwest. Further, the carriers of this haplogroup showed a high proportion of southwestern autosomal ancestry unlike the other N1a1 sublineages. Collectively, these results point to distinct demographics within haplogroup N1a1, possibly induced by two distinct arrival routes into Finland. Overall, our study suggests a more complex genetic population history for Finns than previously proposed.

## INTRODUCTION

Data collected from the Finnish population has been widely used in many genetic studies, ranging from investigations of disease susceptibility to population genetics^1,2^. Relative isolation within the Northeastern corner of Europe (Figure 1), together with small founder populations and several population bottlenecks, have shaped the genetic background of modern Finns distinguishable from other Europeans^3^. Additionally, the Finnish population has been further shaped by various cultural, political, and linguistic influences from differing directions, which have led to a degree of genetic differences seen within the country, most notably between eastern and western Finland^4–7^. These genetic east-west differences, in part illustrated by distribution of Y-chromosomal haplogroups, are suggested to reflect two separate influences from the eastern and western directions^4,7,8^.

**Figure 1.**
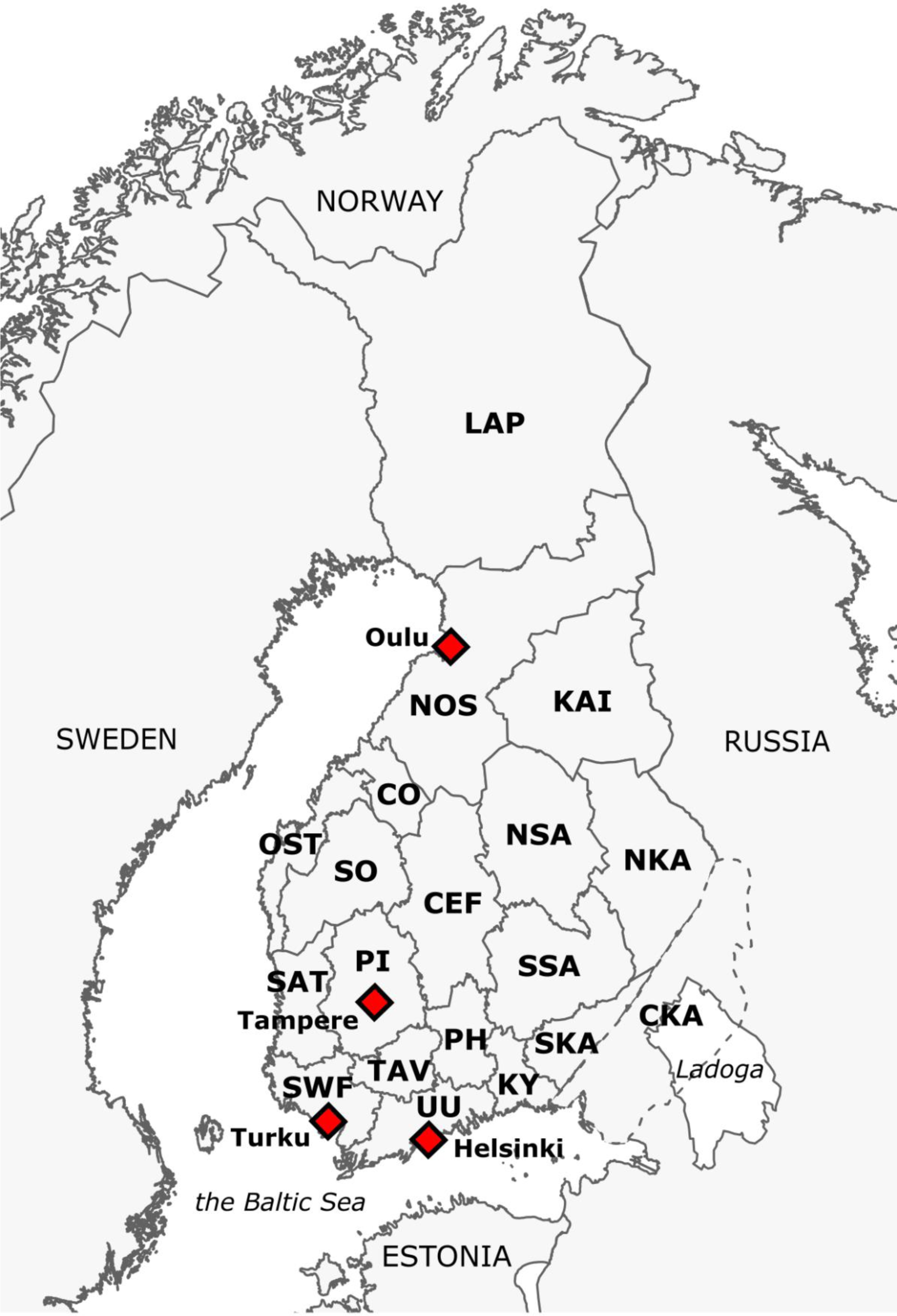
Geographical regions covered in this study. The region names are abbreviated in bold, and their full names are provided in Table 1. Red diamonds highlight 4 of the largest metropolitan areas in Finland. The metropolitan area of Helsinki also includes cities of Espoo and Vantaa. The majority of the Finnish population is located in the southern areas of the country, with Uusimaa (UU) encompassing 31% of the whole population^23^.

**Table 1.** Full names of the studied regions with sample sizes based on the assigned geographical locations. Majority of the samples come from eastern parts of the country due to the sampling strategy of the FINRISK project.

| region | full name | N |
| --- | --- | --- |
| NKA | North Karelia | 392 |
| NOS | Northern Ostrobothnia | 239 |
| NSA | Northern Savonia | 232 |
| LAP | Lapland | 192 |
| UU | Uusimaa | 154 |
| CKA | Ceded Karelia | 111 |
| KAI | Kainuu | 97 |
| SWF | Southwest Finland | 74 |
| SO | South Ostrobothnia | 48 |
| CEF | Central Finland | 48 |
| PI | Pirkanmaa | 46 |
| SSA | Southern Savonia | 40 |
| SKA | South Karelia | 31 |
| KY | Kymenlaakso | 25 |
| SAT | Satakunta | 21 |
| TAV | Tavastia | 16 |
| CO | Central Ostrobothnia | 15 |
| PH | Päijät-Häme | 15 |
| OST | Ostrobothnia | 6 |

The majority of Finnish men belong to the Y-chromosomal haplogroup N1a1 (TAT) (also known as N1c1, N3), having an estimated frequency of 58% in the country^7^. N1a1 represents one of Northeast Eurasia’s prominent patrilineages and is enriched especially among Finno-Ugric populations^9^. Within Finland, the highest frequencies of N1a1 are observed in the eastern regions of the country^7,10^, suggesting eastern introduction of this haplogroup into the country. This aligns with the postulated Siberian origin of the haplogroup^11,12^. The frequency of N1a1 in Europe diminishes rapidly towards the west and south, and it is observed with very low frequencies in Central Europe^13^. Alongside N1a1, a notable proportion of Finnish men belong to haplogroup I1a (M253), carried in total by 28% of men^7^. While I1a is globally enriched in the Scandinavia reaching its peak frequency of 37% in Sweden^14^, unlike N1a1, it is more commonly observed across many European countries^15^. In Finland, I1a is especially frequent along the western coast of the country^7^, aligning with the suggested western influence on this haplogroup^7,10^. In addition to these two major Y-chromosomal lineages, approximately 10% of Finnish men belong to haplogroups R1a (L62) and R1b (CTS2134), which can be associated with Eastern and Western European ancestries, respectively^7^. In Finland, haplogroup R1a has been proposed to have eastern influences via Karelia to the country, whereas R1b has been suggested to have arrived from the western direction^5^.

While previous Y-chromosomal studies in Finland have provided insights into the major substructure of the population, revealing the dual origins of the Finnish gene pool^7,8^, these studies have been limited by the assessment of only a few Y- chromosomal haplogroups determined by genotyping established SNPs and STRs. Recent Y-chromosomal studies within other populations have demonstrated the power of leveraging the combination of sequencing and genotyping data to reveal substructure within major haplogroups, enabling the mapping of these haplogroups into more detailed and time-calibrated population historical events^9^. For instance, Ilumäe et al. (2016), in their comprehensive assessment of haplogroup N1a1 across its entire geographical enrichment area in Northern Eurasia, showed that among northeastern European populations haplogroup N1a1 divides into two distinct sublineages, N1a1a1a1a1a (VL29, CTS2929, N3a3) and N1a1a1a1a2 (Z1936, CTS10082, N3a4), estimated to have become widespread among different parts of the region over the last 5,000 years^9^. Further assessment of such finer level variation within individual populations could potentially provide better detail into the demographics of the Y-chromosomal haplogroups, which further have great potential to elaborate on a population’s history to a deeper detail.

In this study, we characterized the phylogeographic landscape of common Y chromosome variation in Finland by utilizing full sequences of 1,802 Y chromosomes mapped across 19 geographical regions within the country. Overall, our study provides a refined description of the contemporary Y chromosome landscape in Finland, revealing notable heterogeneity especially related to haplogroup N1a1. Overall, our findings suggest that the genetic population history of Finns may be more complex than previously suggested.

## MATERIALS AND METHODS

### Samples

The data for the present study was acquired from the THL biobank (study numbers: BB2019_44, THLBB2022_28) and originated from the FINRISK Project, which is a cross-sectional study of the Finnish working age population with the aim to examine chronic disease risk factors in Finland^16^. FINRISK was initiated in 1972 and it has been carried out in 5-year cycles since its start. Our data set consisted of 1,833 men, whose sex was determined by the registry information, born between 1923 – 1979, included in FINRISK surveys 1992, 1997, 2002 or 2007 with whole-genome sequencing (WGS) data for the Y chromosome available in the biobank (sample sizes by FINRISK surveys presented in Table S1). For each individual, the acquired data included information on their Y chromosome sequence, birthplace, and age. In addition, the majority of the samples had information of their parental birthplaces (N=1,427), autosomal genotyping data (N=1,712), and pre-computed autosomal ancestry profiles (N=758)^17^. These data types are all described in detail later in the methods in their respective sections. All study participants have given a written consent.

### Whole genome sequencing data quality control

Whole-genome sequencing (WGS) was performed for a subset of the FINRISK participants (N=3,322 males and females) at the University of Washington using target coverage of 20x. The reads were mapped to the human genome assembly GRCh38, and variant calling was performed for the whole genome together with thousands of additional samples. Only calls for the Y chromosome were acquired for this project, comprising 1,833 male samples and 295,292 Y-chromosomal variants. Notably, 117,536 (40%) of these sites initial sites were non-polymorphic due to the joint variant calling. We performed variant and sample-wise quality control for the data, removing variants falling on other than X-degenerate, X- transposed or ampliconic regions (definition of MSY regions in Table S2 acquired from^18^), variants without PASS filter, mapping quality ≤20, base quality z-score ≤|2|, strand bias FS >13. All heterozygous calls were set as missing. Sites with >5% missing data, and non-polymorphic sites were excluded, leaving 10,241 variants in the data. Two samples were excluded due to having more than 75% of missing data, leaving 1,831 samples in the data set.

### Allele frequency concordance with Finnish WGS datasets

For the variants passing the quality control, we compared the allele frequency concordance with three WGS datasets of Finns: Sequencing Initiative Suomi (SISu) (N=7,019)^19^, gnomAD v3.1.2 (N=4,029)^20^ and to a combined resource from the 1000 Genomes Project and Human Genome Diversity Project data (1kGP+HGDP) (N=38)^21^. The sequencing data from 1kGP+HGDP was filtered with the same criteria as our FINRISK data, leaving 6,229 variants and 38 Finnish samples in the data. For SISu and gnomAD, which also contain the FINRISK samples used in our study, we only acquired variant level summary data. SISu variants were filtered with call rate ≥95%, heterozygous call rate ≤0.01, removing indels and non-polymorphic variants, yielding 68,624 variants (called for 7,019 Finnish samples). gnomAD variants were filtered to exclude non-polymorphic variants, yielding 44,553 variants (called for 4,029 Finnish samples).

Most of the variants identified in our data were found in these reference data sets (9,756 variants; 95%) with an overall good allele frequency concordance (Figure S1). We accepted the allele frequencies to differ up to 15 percentages between FINRISK and the reference data sets, since the variant frequencies are known to vary extensively based on the geographical location^7^. Out of our variants, 485 (5%) were not found in these reference datasets, with these having low frequencies (MAC ≤ 18 in our data). Given our samples were included also in the reference datasets of SISu and gnomAD, these variants were probably not observed in the reference datasets due differences in filtering steps, thus we decided to remove these 485 variants from our data, leaving 9,756 variants in the final data set.

Overall, most of the detected Y-chromosomal variants were rare (MAF ≤ 0.01) (7,792; 80%), including 3,488 singletons. Out of all variants, 3,348 (34%) were annotated as haplogroup-defining in ISOGG (v15.73) and 1,381 (14%) of the variants were protein coding based on variant effect predictor annotations^22^.

### Geographical location of the samples

Geographical location of the samples was determined by mapping the samples into 19 geographical regions, comprising 18 current administrative regions within Finland and one former Finnish region (Ceded Karelia) that today belongs to Russia (Figure 1). For most of the samples, we utilized the father’s birthplace as the geographical origin (N=1,427), since this enables to map the samples one generation back in time and limits the effects of recent population movements. The remaining samples missing information of their parental birthplaces (N=400), were mapped into the regions using their own birthplace. We removed samples with missing geographical data (N = 4), samples having their birthplace abroad (N = 24), and one individual from Åland Islands due to the low coverage of the region, overall leaving 1,802 samples in the final dataset (Table 1; Figure S2). We acknowledge that the modern administrative regions may not be the most informative regions for population genetic analyses, yet this approach enabled us to preserve individual privacy and still provide sufficient information of the sample’s geographical distribution.

### Haplogrouping and haplogroup nomenclature

Y-chromosomal haplogroups were assigned for each sample with YLineageTracker^24^, which currently is one of the most accurate software tools designed to allocate haplogroups based on VCF formatted data^25^. We annotated the haplogroups and haplogroup-defining variants according to the International Society of Genetic Genealogy (ISOGG) v15.73^26^ nomenclature. Nevertheless, due to the constantly evolving nomenclature system, we also refer to the haplogroups by their defining markers throughout the text.

### Haplogroup frequencies

To assess frequencies for the common haplogroups in Finland with at least 1% frequency, we first selected all terminal haplogroups from YLineageTracker output (i.e., the finest resolution haplogroups that could be classified), and extended this to include also higher nodes within their phylogeny to better group the rare and distinct haplogroups into larger entities. We then filtered these haplogroups to include only common haplogroup-defining variations in the Finnish population, that were observed as a terminal haplogroup or their upper nodes in at least 1% of the data.

The haplogroup frequencies were assessed directly as the allele frequencies of the haplogroup-defining variants, and these are referred to as unscaled frequencies. Since majority of our samples were collected from northeastern parts of the country (Table 1), we further normalized the variant frequencies by weighting each regional frequency estimate with the corresponding population size from Statistics Finland from year 2022^23^, and used these to calculate the frequency estimate within the whole country. This provided more accurate haplogroup frequency estimates among the Finnish population, and these are referred to as scaled frequencies. We further calculated 95% confidence intervals for the frequency estimates of the major haplogroups by the following equation, where p is the haplogroup proportion and n is the total sample size:

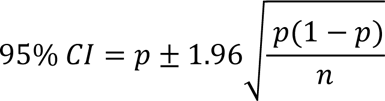

### Phylogeny reconstruction

To assess the phylogenetic relationships of the common haplogroups within the population, we built a maximum likelihood (ML) tree with RAxML v8.2.12^27^. We randomly selected a subset of 55 samples representing distinct haplogroups having at least 1% frequency (haplogroup and its subgroups together) in the data, to focus on the major phylogenetic tree structure and to reduce the computational load of the analysis. We additionally selected one sample with haplogroup E to root the tree. RAxML was ran with the GTRGAMMA model by computing a starting tree from 20 runs, bootstrapping over 100 replicates, and combining these into the final output tree visualized in FigTree v.1.4.3 (Rambaut, 2006-2016) (Figure S3).

### Time estimation

We estimated time to most recent common ancestor (TMRCA) by calculating the average number of newly acquired mutations within the subclades of a given haplogroup node^28^. Since the Y chromosome consists of variable sequence classes having different mutation rates^29,30^, we only considered variation in the X-degenerate region (XDR) within these calculations. Since we were interested here in the ages of haplogroup N1a1 sublineages, we assigned all variants that were polymorphic only within haplogroup N1a1 as “derived” mutations. We then randomly selected one sequence from each of the N1a1 sublineages detected by YLineageTracker (N=98) and calculated the number of derived mutations observed within each haplogroup, and shared mutations within upper nodes according to the phylogenetic tree structure.

Next, we used two methods to derive the TMRCA estimates from the calculated number of mutations. We utilized a calibration point for haplogroup N1a1a1a1a TMRCA at 4995 (4353 – 5700) ya^9^ which yielded with our data a rate of 235,1 (CI: 204,9 – 268,3) years per mutation. Additionally, we used a previously reported mutation rate of 268,5 (CI: 246,3 – 291,9) years per mutation^31,32^ to estimate the TMRCA’s. This mutation rate had been calculated by the rate of 1.0e^-9^ mutations per position per year (CI: 0.92e^−9^ – 1.09e^−9^), generation time of 30 years, and the XDR region length of 3,7 Mb that is uniquely mappable of the XDR^31,33^. Nevertheless, since in our study we only had access to a VCF file we can only assume a similar mapping coverage for the XDR region in our data. Although both these methods rely on many assumptions, e.g., about the mapping quality in our data, mutations occurring at a fixed rate, our estimates were comparable with previous studies^9^.

### Geographical enrichment

To assess the geographical enrichment of the Y-chromosomal variation in the population, we calculated regional frequencies for common haplogroup-defining variations (≥ 1% frequency) and assessed their regional enrichment within the country by a X^2^ test with equal frequencies across all regions as the null hypothesis. We calculated the regional frequencies out of all samples (e.g., I1a out of all samples), and out of major haplogroup carriers (e.g., I1a1a out of I1a carriers), and further visualized both these enrichments on a regional level (online figures). To protect sample privacy and to provide more consistent frequency estimates, on regions with low coverage of samples we used regional averaging when estimating the regional frequencies. This regional averaging was performed by adding all samples from the geographically closest region(s) to the target region until reaching a certain threshold of samples and calculating the frequency estimate using this combined set of samples. When visualizing the enrichments out of all samples, we set a minimum threshold of 15 samples within each region, and out of major haplogroup the threshold was set to at least 10 major haplogroup carriers per region. The visualizations were performed in R utilizing maps from geoBoundaries package^34^. We note that the less frequent haplogroups (e.g., those with 1% frequency in the population) may be inaccurately visualized on a regional level due to the use of regional averaging.

### Autosomal data quality control

To link the Y-chromosomal variation with autosomal genetic variation, we assessed imputed autosomal genotyping data for 1,710 samples. The samples were genotyped with multiple genotyping arrays (Table S1) and imputation was carried out by using the population-specific SISu v3 imputation reference panel with Beagle 4.1 (version 08Jun17.d8b)^35^ as described in the following protocol: dx.doi.org/10.17504/protocols.io.nmndc5e. Before quality control, the dataset consisted of 1,710 individuals and 16,962,023 variants. We performed variant and sample-wise quality control for each chromosome separately in PLINK 2.0^36,37^ removing variants with INFO < 0.99, genotyping quality < 0.99, HW p-value < 1e- 6, sites with > 1% missing data and multiallelic sites. To obtain a set of independent variants, LD-pruning was performed with 1,000 kb windows, step size 1 and with r^2^ threshold of 0.2. After these variant filtering steps, sample-wise quality control was performed by removing individuals if they were born abroad, had missing birth region information, or had excess heterozygosity (deviating more than 4SD units from the mean). After the quality control, the dataset consisted of 1,709 male samples and 119,455 autosomal variants.

### Sample relatedness

Sample relatedness was inferred in PLINK 2.0^36,37^ for the 1,709 samples with autosomal genetic data available. In total 32 sample pairs were classified as closely related (3 pairs as 1^st^ degree, and 29 pairs as 2^nd^ degree), whereas 1,677 samples were classified as unrelated (kinship coefficient < 0.0442). Since 122 samples were lacking autosomal data, we estimated the number of expected relationships that may be present in the whole data. Among our autosomal samples, the rate of 1^st^ degree related pairs was 3/(1709*1708/2) and rate of 2^nd^ degree related pairs was 29/(1709*1708/2), thus we expect that in our whole data of 1,802 we should detect 3.3 of 1^st^ degree and 32.2 of 2^nd^ degree related sample pairs, numbers unlikely to bias the haplogroup frequencies estimated with the whole data. Therefore, we used the whole dataset comprising the 1,802 samples in our main Y-chromosomal analyses, and further utilized the confirmed set of unrelated samples for validating the results. From the unrelated data set we included 1,650 samples in our validation analyses, since these were part of our quality control passing Y-chromosomal dataset having their paternal birthplaces in Finland.

### Principal component analysis

We performed autosomal principal component analysis (PCA) for the 1,709 samples. We first performed PCA in PLINK 2.0^36,37^ for a subset of unrelated samples with autosomal data (N = 1,604) and used the output further in projecting PC scores for all 1,709 samples with autosomal data available. We then used the autosomal PCs to compare their distributions between the carries of different Y-chromosomal haplogroups and performed correlation analysis between the PC scores and Y-chromosomal haplogroup frequencies. The PCs were mapped to geographical regions using the samples’ own birthplaces (Figure S2), since using only the father’s birthplace is an inaccurate measure for autosomal genetic origin.

### Autosomal ancestry profiles by 10 reference populations

To define the autosomal genetic backgrounds of the Y chromosome lineages to a finer and more interpretable detail, we further assessed pre-defined autosomal ancestry profiles created by Kerminen et al. (2021), where the major source of ancestry is accurately detected three generations back in time. The ancestry profiles were based on 10 genetically and geographically mapped Finnish reference populations, and these profiles were available for 758 samples in our data. We assessed the major source of ancestry for each sample by the criteria of sharing at least 50% of their genome with one reference population, resulting in 485 samples with one major source of ancestry. We used these autosomal ancestry profiles to compare their distributions between different Y- chromosomal haplogroup carriers.

## RESULTS

### Y-chromosomal variation in Finland

To characterize common Y-chromosomal variation in Finland, we analyzed 1,802 geographically mapped high-coverage Y chromosome sequences obtained from the FINRISK project^16^. Within this dataset we identified a total of 111 distinct haplogroup-defining variants observed with at least 1% in the population (Table S3; Figure 2A-B). Notably these variants were not exclusively terminal haplogroups (i.e., the finest resolution haplogroup that could be classified), but also included internal branches of the haplogroup tree. Removing the internal branches, we used 55 of the total 111 common haplogroups (i.e., assigned as terminal haplogroup for at least one sample) for visualizing the main relationships and clustering of Finnish Y chromosomes in a maximum-likelihood phylogenetic tree (Figure 2C).

**Figure 2.**
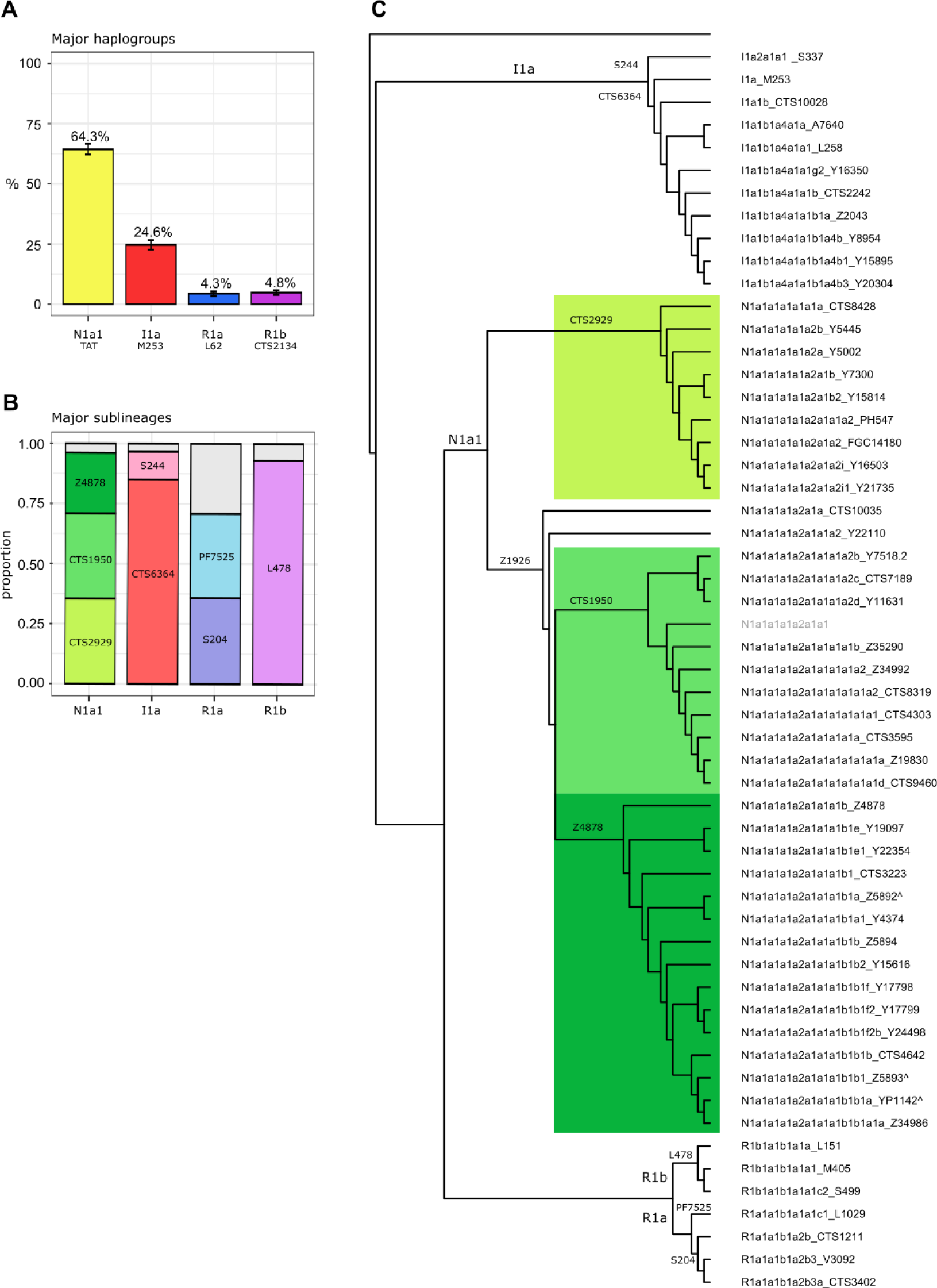
Common Y-chromosomal variation in Finland. A) Frequencies of the four major haplogroups, B) proportions of the most common sublineages detected among these, and C) the phylogenetic relationships and subclustering of detailed sublineages. The frequencies indicated on the figure correspond to scaled frequencies (i.e., normalized by regional population sizes). The phylogenetic tree is not scaled to time, with the branch lengths being proportional to the number of tips under the node. Haplogroup N1a1a1a1a2a1a1 marked in gray has an ambiguous position in the tree, possibly indicating a more detailed haplogroup for this sample than YLineageTracker could classify.

We identified the four previously described main haplogroups in Finland (N1a1, I1a, R1a, and R1b) (Figure 2A), with similar frequency estimates as previously described^7,10^. However, with sufficient sample coverage across the country and through the scaling of our estimates by regional population sizes (see Methods), our data enabled us to provide refined frequency estimates for these main haplogroups. Among the Finnish population, haplogroup N1a1 (TAT) accounted for 64.3% (95% CI 62.1 – 66.5%), I1a (M253) for 24.6% (95% CI 22.6 – 26.6%), R1a (L62) for 4.3% (95% CI 3.4 – 5.2%) and R1b (CTS2134) for 4.8% (95% CI 3.8 – 5.8%) of the Y chromosomes (Figure 2A; Table S3). The remaining 2% of samples carried haplogroups previously recognized as rare in the Finnish population^7,10^.

Importantly, our data allowed for identification of several sublineages within these previously described major haplogroups (Figure 2B-C; Table S3). The most notable observation was the subdivision of haplogroup N1a1 into three major sublineages within Finland (Figure 2B-C). Practically all N1a1 carriers belonged to the haplogroup N1a1a1a1a (F4155) after which the haplogroup divided into sublineages N1a1a1a1a1a (CTS2929) (35.5% of N1a1), N1a1a1a1a2a1a1a1a (CTS1950) (35.4% of N1a1) and N1a1a1a1a2a1a1a1b (Z4878) (25.2% of N1a1) (Figure 2B-C; Table S3). We estimated the TMRCAs for haplogroup N1a1a1a1a1a (CTS2929) at 4,995 ya, for N1a1a1a1a2a1a1a1a (CTS1950) at 1,754 ya and for N1a1a1a1a2a1a1a1b (Z4878) at 2,986 ya, respectively (Table S4).

Within haplogroup I1a we observed several sublineages, most notably splitting into haplogroups I1a1 (CTS6364) (20.8%) and I1a2 (S244) (3.3%) (Figure 2B; Table S3), as previously described^10^. The most common sublineage within haplogroup I1a was I1a1b1a4a1a1 (L258) (16.8%), which further divided into lineages I1a1b1a4a1a1b (CTS2242) (6.0%) and I1a1b1a4a1a1g (Y15027) (2.3%) (Table S3).

Within haplogroup R1a we found that all samples belonged to haplogroup R1a1a1b (PF6158), after which it dived into two major sublineages: R1a1a1b1a1a (PF7525) (1.5%) and R1a1a1b1a2 (S204) (1.7%) (Figure 2B; Table S3). Within haplogroup R1b all samples belonged more specifically to R1b1a1b1 (L478) and its sublineages (Figure 2B; Table S3).

### Geographical distribution of Y-chromosomal haplogroups in Finland

Previous work have identified differences in the geographical distribution of haplogroups N1a1 and I1, especially between the eastern and the western parts of Finland, suggested to reflect two distinct migration directions into the country^7,10^. Nevertheless, with a sample size three times greater than in previous studies and a more comprehensive sampling across the in the country, we were able to comprehensively reassess the major haplogroup distribution (Figure 3A), and further extend this beyond the major haplogroups to all 111 common haplogroup-defining variants within the country (Table S5; Table S6; online figures). We assessed the geographical distribution on two levels, by the regional frequencies corresponding 1) to each haplogroup’s proportion of all samples (Table S5) and 2) to each haplogroup’s proportion of its major haplogroup (N1a1, I1a, R1a or R1b) (Table S6).

**Figure 3.**
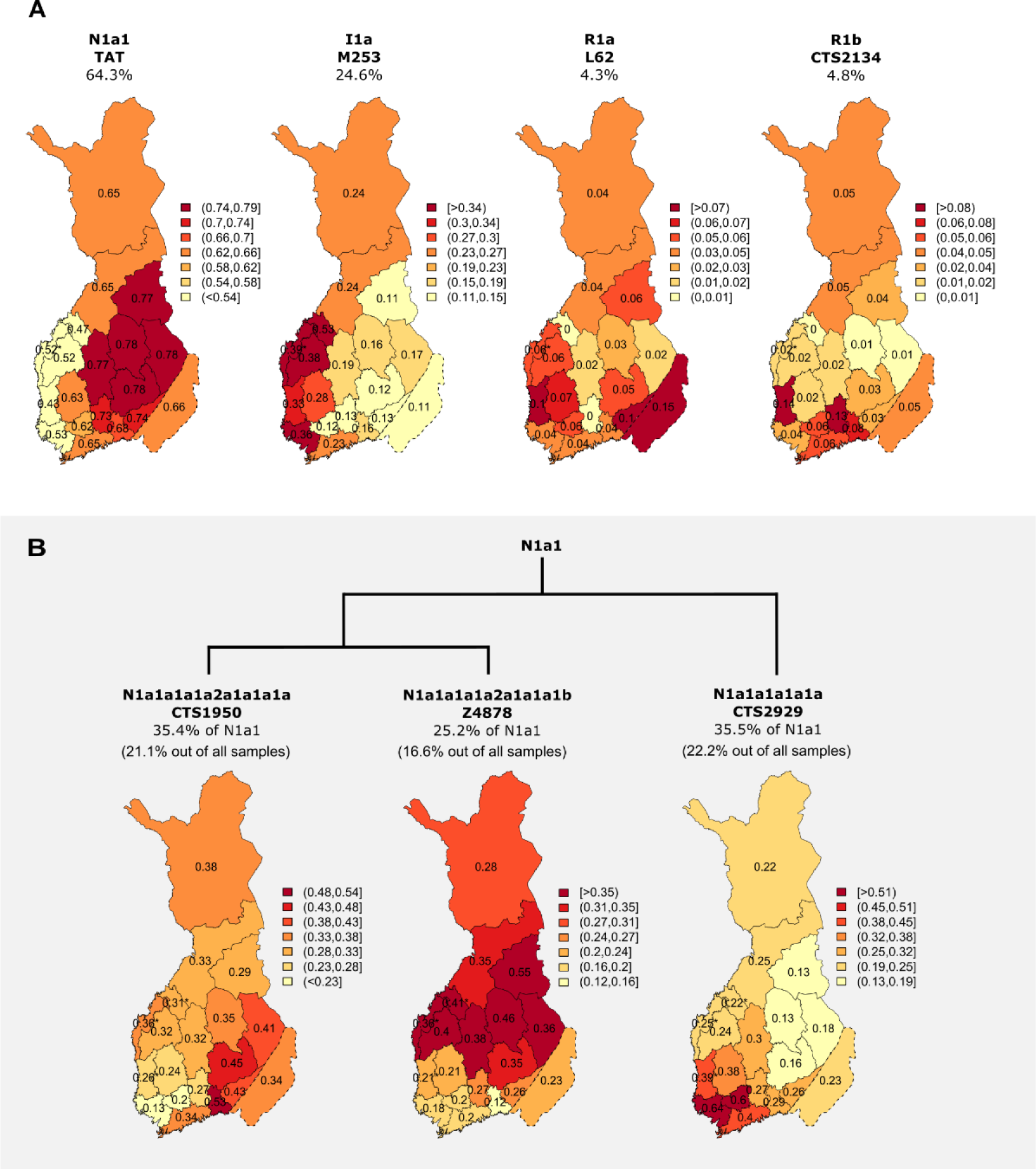
Geographical distribution of A) major Y-chromosomal haplogroups N1a1, I1, R1a, R1b, and B) the three major N1a1 sublineages within Finland. The coloring indicates haplogroup frequency and is scaled for each map separately with the average frequency as the midpoint. In panel A the haplogroup frequencies were calculated out of all samples, whereas in B the haplogroup frequencies are calculated out of the major haplogroup N1a1. * = haplogroup frequency is imputed from geographically closest regions due to low coverage of samples in the region. In panel A, the frequency for the region OST is imputed, and within panel B regions OST, CO and SAT the frequencies are imputed. The frequencies indicated within the figure headers correspond to scaled frequencies.

### Major haplogroups N1a1, I1a and R1a show east-west differences

Assessing the geographical enrichment of the 111 haplogroup-defining variants, we observed in total 65 lineages displaying nominal geographical enrichment (p < 0.05 within the whole dataset and the unrelated subset) (Table S5). As previously reported^7,10^, we observed strong geographical enrichment for haplogroups N1a1 and I1a (Figure 3A; Table S5). Haplogroup N1a1 reached its highest frequency of 78% in North Karelia, North Savonia, and Southern Savonia (p = 3.1×10^-7^, X^2^ test for equal proportions, df=18), corresponding to a 1.2-fold enrichment compared to the haplogroup frequency within the country (Figure 3A; Table S5). In contrast, haplogroup I1a displayed an enrichment along the western coast of Finland, reaching its peak frequency of 53% in Central Ostrobothnia corresponding to a 2.2-fold enrichment (p = 6.6×10^-7^, df = 18) (Figure 3A; Table S5). In addition to these previously established findings, we discovered further heterogeneity in the geographical distribution of haplogroup R1a (p = 1.2×10^-3^, df = 18), displaying a dual enrichment in the east (15% in Ceded Karelia) and in the west (10% in Satakunta) (Figure 3A). Although haplogroup R1b also initially displayed geographical enrichment (p = 0.017, df = 18) reaching its highest frequency in Satakunta (Figure 3A), this result, however, did not replicate in a subset of unrelated individuals (p = 0.22, df = 18) (Table S5), plausibly implying a more dispersed enrichment pattern for R1b throughout the country, or could be related to the overall small number of R1b haplogroup carriers (N=64) in our data.

### Substantial regional heterogeneity beyond the major haplogroups

We further compared the subsequent sublineages of each of the major haplogroups (N1a1, I1a, R1a, R1b) to assess their distributions in better detail without being impacted by the major haplogroup enrichment pattern. With this approach we observed 38 sublineages showing regional heterogeneity (p < 0.05), with 19 of these remaining significant after multiple testing correction (p < 0.05/86) (Table S6). Overall, the majority of these geographically enriched haplogroups were related to haplogroup N1a1 sublineages.

Out of the three identified N1a1 sublineages (Figure 2B-C), haplogroups N1a1a1a1a2a1a1a1a (CTS1950) (35% of N1a1) and N1a1a1a1a2a1a1a1b (Z4878) (25% of N1a1) displayed enrichment predominantly to southeast and northeast of the country, respectively (Figure 3B, Table S6), with these distributions being fairly expected given the N1a1 geographical enrichment pattern in the east (Figure 3A). While N1a1a1a1a2a1a1a1b (Z4878) reached its highest frequency of 55% of N1a1 in Kainuu (p = 7.5×10^-6^, df = 18), haplogroup N1a1a1a1a2a1a1a1a (CTS1950) reached its highest frequency in Kymenlaakso, although this enrichment was not statistically significant (Figure 3B; Table S6). In contrast to these eastern enriched N1a1 lineages, haplogroup N1a1a1a1a1a (CTS2929) (36% out of N1a1) was enriched to the opposite side of the country (p = 3.8e^-11^, df = 18) (Figure 3B; Table S6). N1a1a1a1a1a (CTS2929) reached its highest frequency of 64% of N1a1 in Southwest Finland (Figure 3B). This lineage further dived into two distinct sublineages N1a1a1a1a1a1 (Z4908) (14%) and N1a1a1a1a1a2 (CTS9976) (22%), with slightly differing enrichment patterns from each other (Table S6; online figures), nevertheless both being clearly enriched to the southwest (Table S6).

Within haplogroup I1a, the majority of its sublineages exhibited the expected enrichment into the western regions similarly to the major haplogroup (Table S6). Some of the I1a sublineages further displayed enrichment to the east, such as I1a1b1a4a1a1b1a4 (Y10990) (13.2% of I1a) reaching its highest frequency of 42% of I1a in Ceded Karelia (p = 5.0×10^-4^, df = 18). However, for haplogroup R1a and R1b sublineages, we could not find enrichments beyond the major haplogroup level that would suggest a clear centralized area of enrichment, although haplogroup R1b1a1b1a1a (R-L151) (93% of R1b) displayed a nominal evidence for heterogeneity across the regions (p = 0.012, df = 18) (Table S6). The lack of further observed enrichment patterns within the R1a and R1b sublineages may have been impacted by to their lower frequencies in the data, we nevertheless observed a relatively even distribution for them throughout the country.

### Autosomal genetic structure correlates with major haplogroups N1a1 and I1a in Finland

In addition to the Y-chromosomal haplogroups, the autosomal genetic structure is known to vary geographically in Finland, with the largest differences observed between eastern and western parts of the country^6^ (Figure 4A). To gain insights into this connection between the autosomal genetic population structure and Y- chromosomal haplogroups within the country, potentially providing demographic insights for the Y-chromosomal haplogroups, we extended our analyses to examining autosomal genetic structure. To this end, we compared the autosomal genetic background between carriers of different Y-chromosomal haplogroups using two measures, 1) autosomal PCs 1-20, typically used in the characterization and adjustment of genetic population structure (Figure 4A), and 2) previously described autosomal ancestry profiles from Kerminen et al. (2021), at the level of 10 reference populations representing the Finnish population structure in a finer and more interpretable detail (Figure 4F).

**Figure 4.**
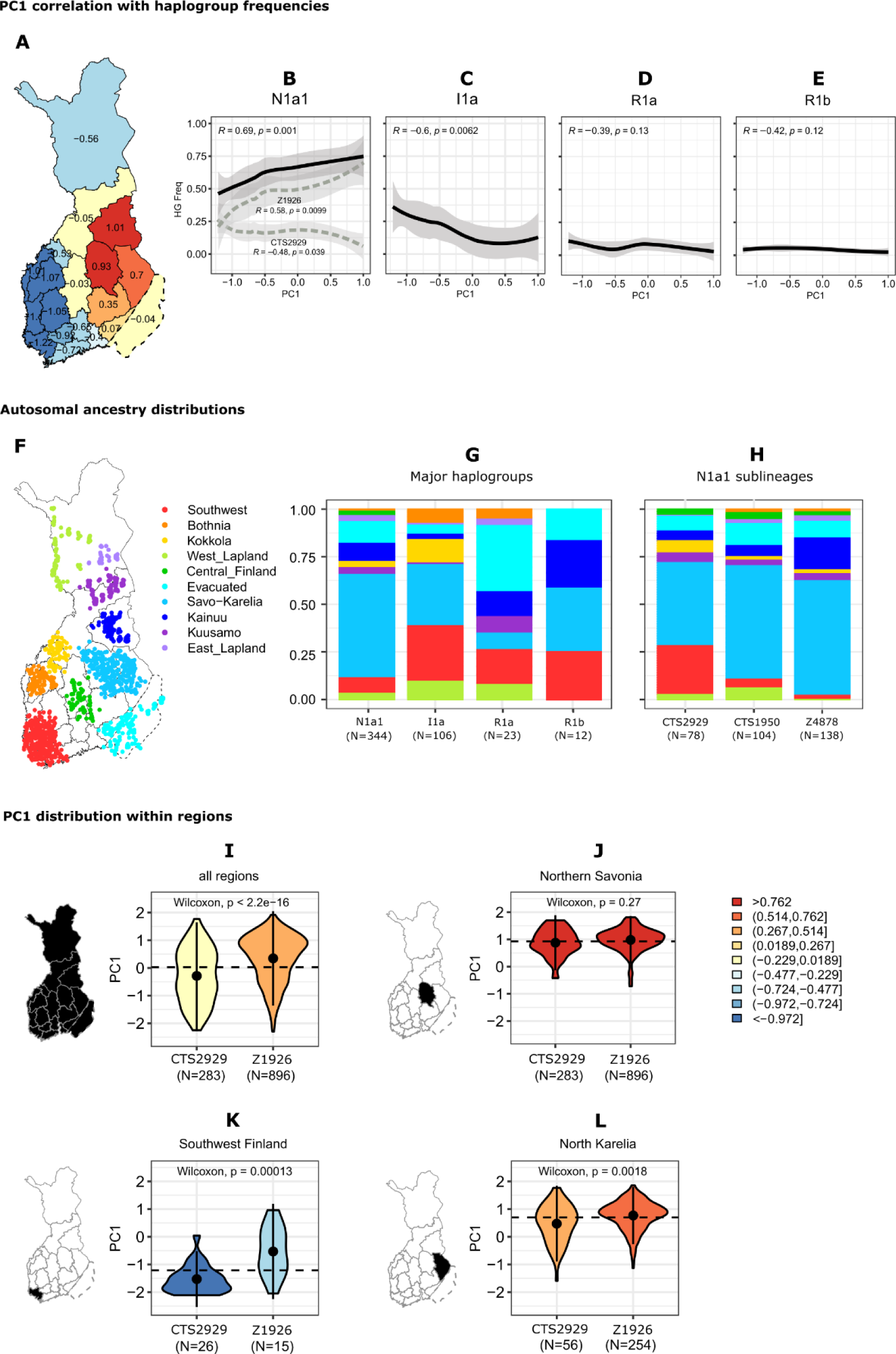
Connection between the autosomal genome and Y chromosomal haplogroups. A-E) The correlation between autosomal PC1 and Y-chromosomal haplogroup frequencies regionally for the major haplogroups. The lines are visualized by LOESS. F-H) Autosomal ancestry distributions by 10 FineSTRUCTURE reference populations from Kerminen et al. (2021), with the coloring ordered by the populations’ approximate Fst distances. Each individual was assigned to one population if sharing at least 50% of their genome with the reference population. I-L) Regional distribution of autosomal PC1 regionally compared between haplogroups N1a1a1a1a1a (CTS2929) and N1a1a1a1a2a1a1 (Z1926). The coloring in panels I-L corresponds to autosomal PC1 scores.

PC1 captures the geographically varying east-west autosomal genetic substructure among Finns (Figure 4A), with this pattern being similar to several Y-chromosomal haplogroup distributions which were most notably varying along the east-west axis (Figure 3A; Table S5; Table S6). To quantify this relationship further, we calculated the correlation between regional haplogroup frequencies and regional PC1 scores within the country. At the major haplogroup level, we observed a significant correlation between PC1 and haplogroups N1a1 (R = 0.69, p = 0.001) and I1a (R = -0.60, p = 0.006) (Figure 4B-C). In contrast haplogroups R1a and R1b did not show significant correlation with the PC1, likely impacted by their more dispersed enrichment patterns.

When further assessing the PC1 correlation within the N1a1 subgroups, we observed that the PC1 correlation pattern of the southwestern enriched lineage N1a1a1a1a1a (CTS2929) was opposite to that of the main haplogroup N1a1 (R = -0.47, p = 0.043) (Figure 4B). This observation suggests that the correlation between the autosomal PCs and Y-chromosomal haplogroups captured on the major haplogroup level is not necessarily representative of the relationship within the further sublineages.

As a complementary approach, we employed autosomal ancestry profiles from 10 reference populations acquired from Kerminen et al. (2021) (Figure 4F), which offer a finer and more interpretable representation of the population structure compared to PCs alone. Here, each sample was assigned with an ancestry based on the criteria of sharing at least 50% of the genome with one of the reference populations, leaving 485 samples with one major source of ancestry for the analyses. Subsequently, we examined the distributions of these autosomal ancestry assignments among carriers of different haplogroups (Figure 4G-H). For the major haplogroups, we identified expected differences in their autosomal ancestry distributions (Figure 4G; Figure S4). Most of haplogroup N1a1 carriers belonged to the eastern Savo-Karelia ancestry, while carriers of haplogroup I1a displayed higher proportions of western ancestries, such as Southwest, Bothnia, Kokkola, and West Lapland compared to N1a1 (Figure 4G; Figure S4). Within haplogroup R1a we observed the highest proportion of Evacuated ancestry, corresponding to the region of Ceded Karelia (Figure 4G; Figure S4). While R1b displayed visually higher proportions of Kainuu ancestry (Figure 4G), the proportion was not significantly higher than for R1b or N1a1 (Figure S4), likely impacted by the small sample size for R1b (N=12) in the analyses.

### N1a1 sublineage carriers show distinct autosomal genetic ancestry

To further investigate whether we could observe differences within the autosomal genetic background of the major N1a1 sublineages, indicative of distinct recent demographics, we conducted a comparison of autosomal ancestry proportions for carriers of the three N1a1 sublineages: N1a1a1a1a1a (CTS2929), N1a1a1a1a2a1a1a1a (CTS1950) and N1a1a1a1a2a1a1a1b (Z4878) (Figure 4H; Figure S5). The northeastern enriched lineages, N1a1a1a1a2a1a1a1a (CTS1950) and N1a1a1a1a2a1a1a1b (Z4878), were both enriched in Savo-Karelia ancestry in a similar manner to the main haplogroup N1a1 (Figure 4G). However, carriers of the southwestern enriched lineage N1a1a1a1a1a (CTS2929) deviated from this ancestry pattern, displaying a higher proportion of southwestern ancestry compared to the other N1a1 lineages (Figure 4H; Figure S5).

### Regional differences in PC1 indicate recent population movements

To further investigate a possible southwestern origin for haplogroup N1a1a1a1a1a (CTS2929), we compared its autosomal genetic background to N1a1a1a1a2a1a1 (Z1926) within individual regions. To this end, we used the autosomal PC1 scores and compared their distributions between these lineages. Within Southwest Finland, we observed significant differences between the PCs for the carriers of the southwestern N1a1a1a1a1a (CTS2929) (N = 29) and northeastern N1a1a1a1a2a1a1 (Z1926) (N = 15) haplogroups (Wilcoxon p = 1.3×10^-4^) (Figure 4K), suggesting distinct demographics for these two haplogroups within this region. Carriers of the southwestern lineage N1a1a1a1a1a (CTS2929) were enriched towards lower PC1 values (typical for samples of southwestern origin), whereas carriers of the northeastern lineage N1a1a1a1a2a1a1 (Z1926) were enriched towards higher PC1 values (typical for northeastern regions). We also observed autosomal differences for these haplogroups in North Karelia, where similarly carriers of the southwestern N1a1a1a1a1a (CTS2929) were enriched towards lower PC1 values compared to the northeastern N1a1a1a1a2a1a1 (Z1926) (Wilcoxon p = 1.8×10^-3^) (Figure 4L). Altogether these findings highlight the distinct autosomal genetic characteristics within these haplogroup N1a1 sublineages, supporting the idea of a possible southwestern introduction for haplogroup N1a1a1a1a1a (CTS2929) and eastern introduction for N1a1a1a1a2a1a1 (Z1926) into the country.

## DISCUSSION

While the autosomal genome provides a good description of the contemporary genetic structure of a population, the paternally inherited Y chromosome has great potential to elaborate on the population history due to its genetic material not getting recombined through generations. Previous studies have extensively characterized the fine-scale population substructure within Finland, focusing on the autosomal genome^6,17^. However, Y-chromosomal genetic variation among the Finns has remained relatively coarsely characterized, at the resolution of a few genetic markers across a limited number of geographical areas^7,10^.

In this study, we set out to study the Y-chromosomal landscape in Finland by assessing 1,802 Finnish Y chromosome sequences from the FINRISK project. Our data consisted of Finnish men born between 1923 – 1979 with high geographic coverage among the country. Since we used paternal birthplaces for geographical mapping of the samples, our data reflects the Y-chromosomal landscape from the beginning of the 20^th^ century to approximately the 1950’s, before the start of the large-scale internal movements and urbanization within Finland. Employing the combination of high-coverage sequencing data and extensive geographical coverage, together with a large sample size, allowed for a detailed exploration of Y-chromosomal variation within Finland. The resolution of our data enabled the subdivision of previously described major haplogroups (N1a1, I1a, R1a, R1b) in Finland into numerous sublineages common in the population and uncovering novel geographical heterogeneity within them. Overall, our findings suggest more complex composition in the paternal lineages in Finland than previously thought, suggesting, for instance, a dual entry for haplogroup N1a1 into the country.

Previously, haplogroup N1a1 (TAT) has been identified as a prominent patrilineage among Finns^7,10^, carried by 64% of Finnish men according to our estimate. Its prevalence is particularly pronounced in eastern Finland, aligning with a proposed eastern influence into Finland within the last millennia^7,10^. Confirming this previously reported eastern enrichment for N1a1 within Finland, our data highlights the specific contribution of haplogroup N1a1a1a1a2a1a1a (Z1926) driving this enrichment pattern. This haplogroup is carried by 63% of all N1a1 carriers, with an overall frequency of 42% among Finnish men. Globally, N1a1a1a1a2a1a1a (Z1926) is known to show high frequency among Finns, reaching notable frequencies also in the neighboring regions towards the east, e.g., among Vepsas, Karelians, Saamis, and North Russians^38^. Furthermore, the lineage N1a1a1a1a2a1a1a (Z1926) descends from N1a1a1a1a2 (Z1936, CTS10082), a haplogroup which has been associated as a plausible connection among members of the Finno-Uralic language family^39^. Within Finland, haplogroup N1a1a1a1a2a1a1a (Z1926) further divides into two main lineages, with a “Savonian” sublineage N1a1a1a1a2a1a1a1b (Z4878) enriched in the northeast, and a “Karelian” sublineage N1a1a1a1a2a1a1a1a (CTS1950) displaying a more dispersed enrichment pattern in the southeast. A possible source of haplogroup N1a1a1a1a2a1a1a (Z1926) into the country could be through migrations from Siberia^12^ that started to arrive in Northeastern Europe around 3,500 years ago^40^. According to our estimate the “Savonian” and “Karelian” sublineages share a common ancestor approximately 3,200 years ago, which could indicate the split of these two groups occurred in the close proximity of Finland. The presence of many sublineages for these two haplogroups dispersed throughout the country might reflect population expansion events during the late settlement process of Finland over the past millennium^41^.

In addition to the eastern enriched lineages of N1a1, approximately one third of N1a1 carriers belong to haplogroup N1a1a1a1a1a (CTS2929), overall carried by 22% of Finnish men. Globally, recognized as the Baltic branch of N1a1, this haplogroup reaches its highest frequencies among Estonians (28%), and is further present among Latvians, Lithuanians, Finns, Saami, Karelians, Belarusians, Ukrainians, Russians^9^. Within Finland, this haplogroup displays strong geographical enrichment to the southwestern coast of Finland, with the haplogroup carriers also exhibiting a high proportion of southwestern autosomal genetic ancestry. This geographical distribution pattern and the autosomal genetic background of haplogroup N1a1a1a1a1a (CTS2929) carriers distinguish the lineage from the other N1a1 sublineages within Finland. Overall, observing N1a1a1a1a1a (CTS2929) in high frequencies in the southwest (64% of N1a1) and in low frequencies in the east (18% of N1a1) contradicts with the suggested solely eastern route of N1a1 into the country.

Collectively, these findings indicate the potential source of haplogroup N1a1a1a1a1a (CTS2929) in the southwestern regions of Finland is across the Baltic Sea, potentially originating from Estonia where the haplogroup is frequent^9^. In addition to Estonia being the nearest country to Finland across the Baltic Sea, the two populations share close historical, linguistic, and genetic connections with each other. Southwest Finland in particular, with its coastal location along the Baltic Sea, could have been influenced by such gene flow. Nowadays this area contains one of the largest, and also the oldest city of Finland, Turku. Additionally, the Southwestern Finnish dialect spoken within the area stands out for its similar features to the Estonian language in comparison to other Finnish dialects^42^. Such a potential genetic influence from Estonia could originate from a distant migratory event, or alternatively from a more recent event such as the late migration from Estonia to Finland around 1,300 to 1,100 years ago^43^. However, we cannot determine the arrival time of the haplogroup into the country by utilizing only contemporary DNA. Furthermore, since the haplogroup has also been observed among Swedes^44,45^, although with a low frequency (4.4%)^46^, we nevertheless cannot exclude a more complex pattern of migration affecting the enrichment of N1a1a1a1a1a (CTS2929) in Southwest Finland based on the data of this study.

Beyond haplogroup N1a1, the Finnish population is further enriched in haplogroup I1a (M253), which is carried by 25% of Finnish men according to our estimate. As reported previously, haplogroup I1a reaches its highest frequencies along the western coast of Finland, in concordance with the suggested Scandinavian influence of this haplogroup into the country^7^. While we find a predominantly western enrichment for the majority of the haplogroup I1a lineages, we further distinguish an eastern enrichment for sublineage I1a1b1a4a1a1b1a4b (Y8954), carried by 2% of the population. While this enrichment pattern could indicate an eastern direction of arrival, the gradual shift from west to east in the enrichment pattern seen for the phylogenetically higher nodes of this lineage (such as I1a1b1a4a1a1b, CTS2242), rather suggests population movements from the west causing this eastern enrichment pattern.

In addition to haplogroups N1a1 and I1a carried by the vast majority of Finnish men, approximately 10% of Finnish men belong to haplogroups R1a (L62) and R1b (CTS2134)^7,10^. While haplogroup R1a (carried by 4.3% of Finnish men) has a speculated influence from the eastern direction, haplogroup R1b (carried by 4.8% of Finnish men) has been suggested to arrive from the west^47^, aligning with the global enrichment patterns of these two haplogroups^48,49^. We find that haplogroup R1a is geographically enriched in the east but is also observed in high frequencies locally in the west. While this enrichment pattern could indicate a dual influence of R1a into the country, the fact that in our data we primarily identified R1a sublineages that have previously been reported among Russians and Balts, but not in Swedes^50^, support a major eastern influence for R1a into the country. In contrary, for haplogroup R1b we did not find any significant enrichment, implying a relatively equal spread for it across the country. Nevertheless, the lack of the any observed enrichments could partly be influenced by the relatively small sample size for this haplogroup within our data (N = 64).

In summary, we provide a comprehensive exploration of Y-chromosomal variation in Finland, moving beyond a few major haplogroup lineages to unraveling finer-scale variation in the population. In addition to detecting extensive variation in the Finnish Y-chromosomal haplogroups, we further describe geographical heterogeneity among these lineages, in particular related to haplogroup N1a1. Observing geographical differences within the major lineages of haplogroup N1a1, and further differing autosomal genetic backgrounds within the carriers, overall suggest distinct demographics within haplogroup N1a1. We suggest haplogroup N1a1 most likely arrived via two distinct routes to the country, with the major influence rising from the northeast via the mainland, and a subsequent influence from the southwestern direction via the Baltic Sea. Overall, our results highlight that studying the paternally inherited Y chromosome using WGS data mapped to precise geographical origins, has potential to capture additional population historical events compared to autosomal genetic data alone.

## DATA AVAILABILITY

The data used in this study is available through the National Institute for Health and Welfare Biobank (http://www.thl.fi/biobank). Online figures include regional enrichment maps for all common Y-chromosomal haplogroup-defining variants (link to be provided). Supplementary material includes Figures S1-S5 and Tables S1-S6.

This is an open-access article distributed under the terms of the Creative Commons Attribution 4.0 International License (https://creativecommons.org/licenses/by/4.0/), which permits unrestricted use, distribution, and reproduction in any medium, provided the original work is properly cited.

## ACKNOWLEDGEMENTS

The data used for the research was obtained from THL Biobank (study numbers: BB2019_44, THLBB2022_28). We thank all study participants for their generous participation in biobank research. We thank Priit Palta and Shuang Luo for providing the SISu refence data for the Y chromosome. We thank Elina Salmela for insightful discussions and comments on the manuscript.

## AUTHOR CONTRIBUTIONS

T.T., J.L., A.P., and M.P. designed the study. A.P. conducted the analyses, and J.R. provided assistance and computational methods in the analyses of autosomal data. M.P. provided materials and statistical assistance. A.P, J.L., and T.T. wrote the manuscript. All authors interpreted the results and reviewed the manuscript.

## ETHICAL APPROVAL

The data used in this study originated from the FINRISK study, for which ethical approval had been obtained at the time of each survey according to the Finnish legislation and common ethical requirements^16^.

