## Supplementary Figures S1-S5 for "Y chromosome sequencing data suggests dual paths of haplogroup N1a1 into Finland"

**Figure S1**

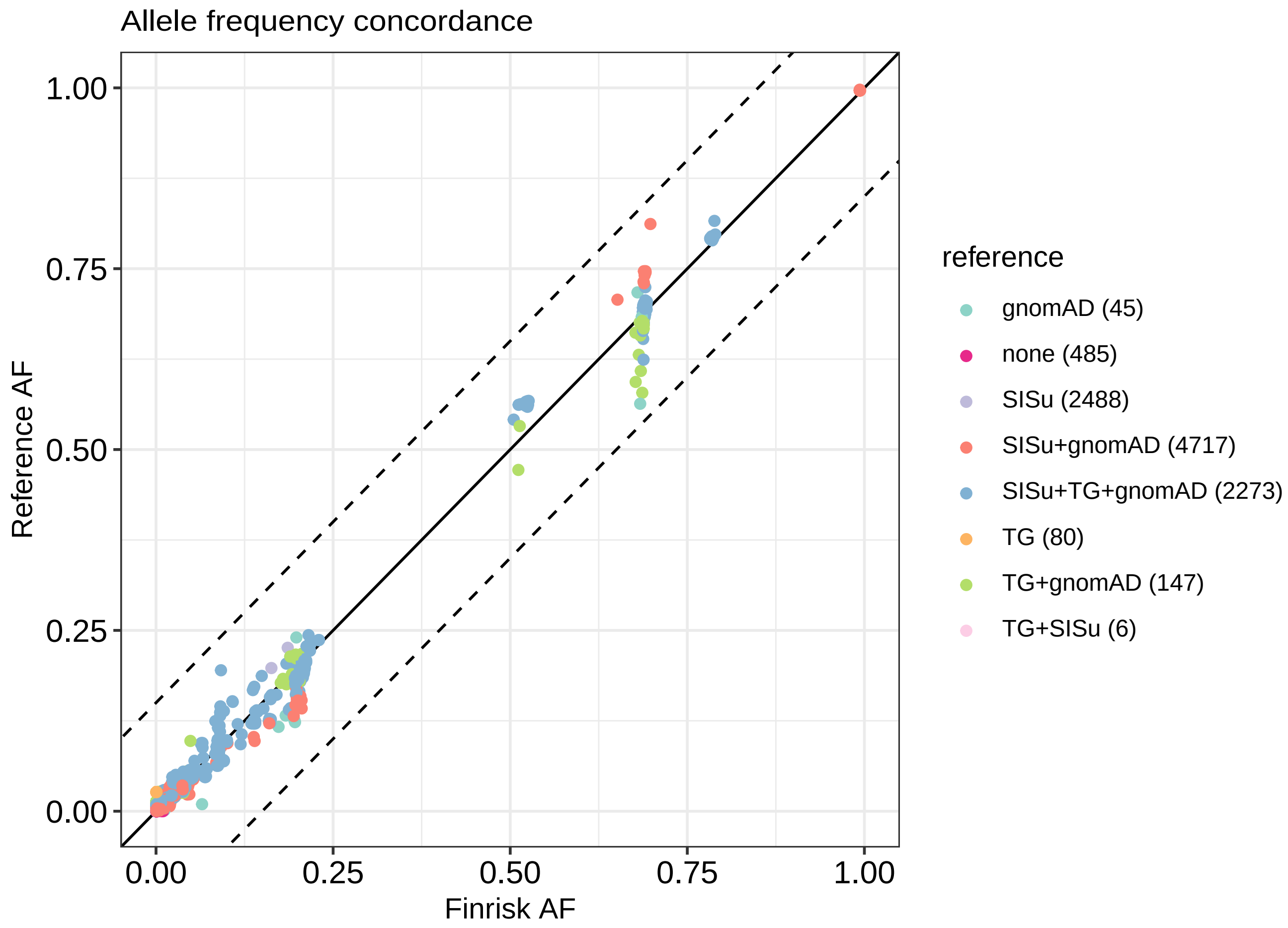


Figure S1: Allele frequency concordance with three Finnish WGS datasets: SISu, 1000 Genomes Project Finns (TG), and gnomAD v3.1.2 Finns.

**Figure S2**

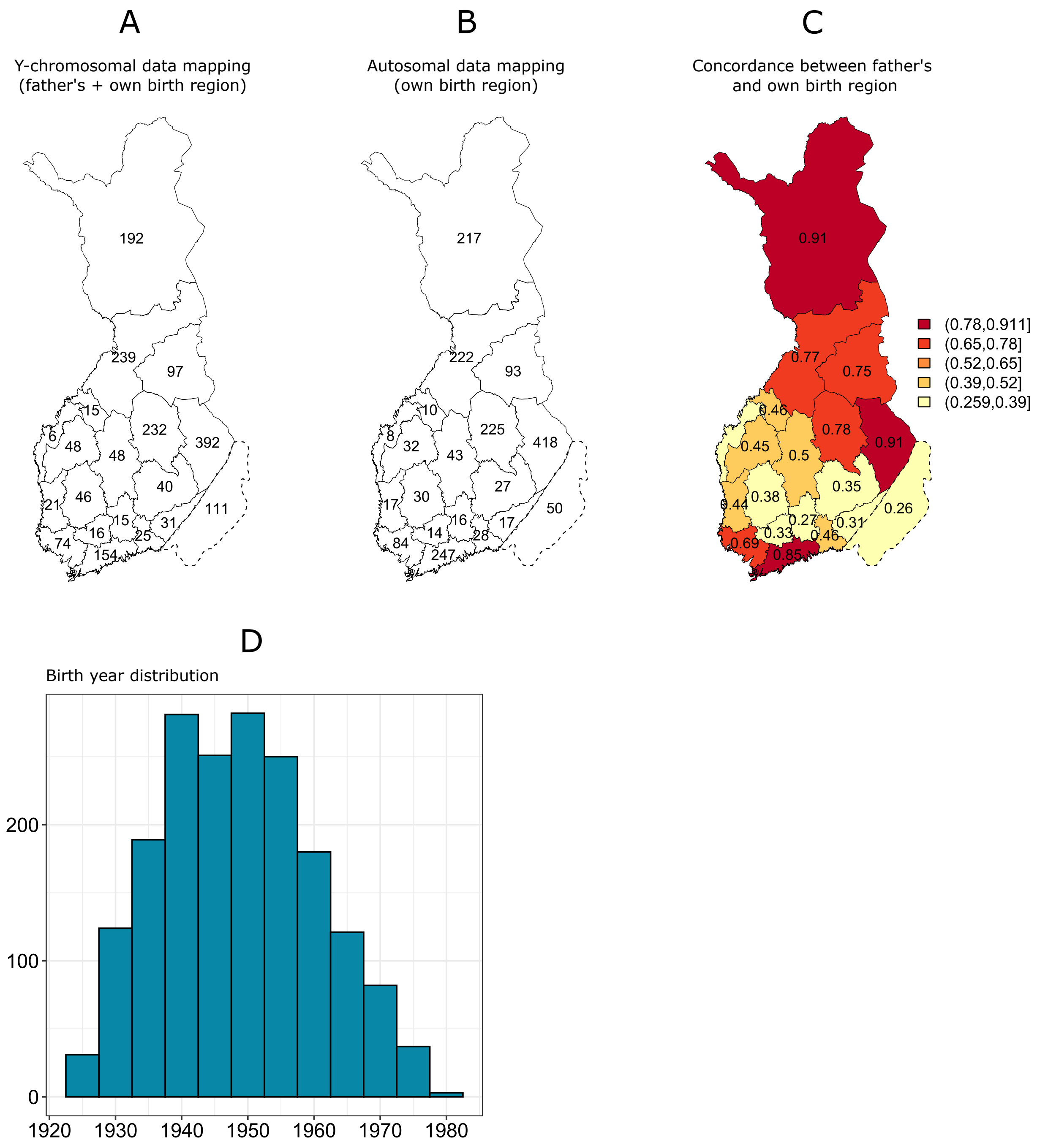


Figure S2: A) Sample distribution for Y-chromosomal analyses (N=1,802), and B) autosomal analyses (N=1,709). C) Concordance between father's and own birth region indicated as a proportion of fathers having their son born in the same region. D) Sample distribution by birth year.

### Figure S3

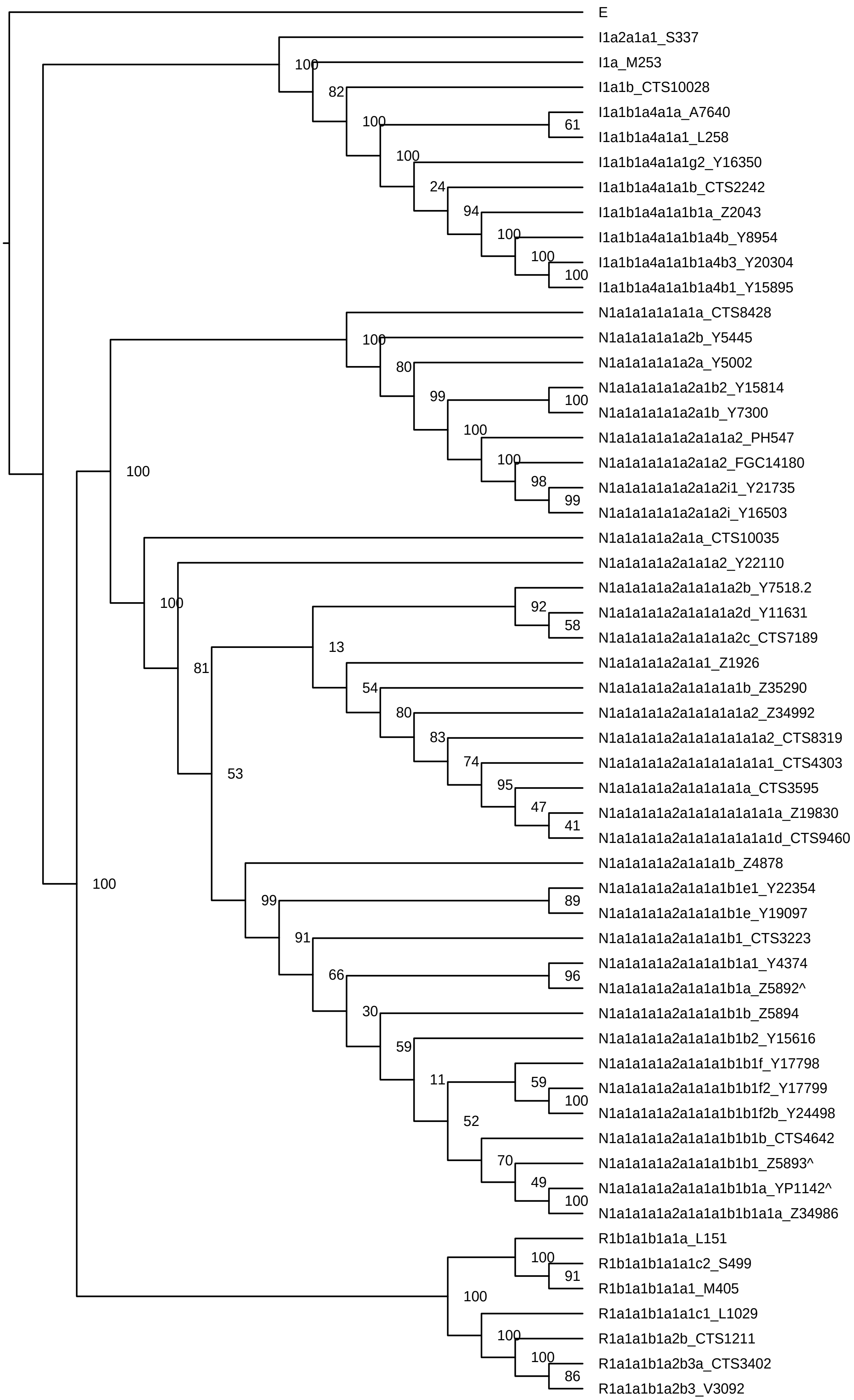


Figure S3: Phylogenetic tree including bootstrap values with subset of 55 samples not scaled to time. Haplogroup label according to ISOGG v15.73 and terminal marker ID.

**Figure S4**

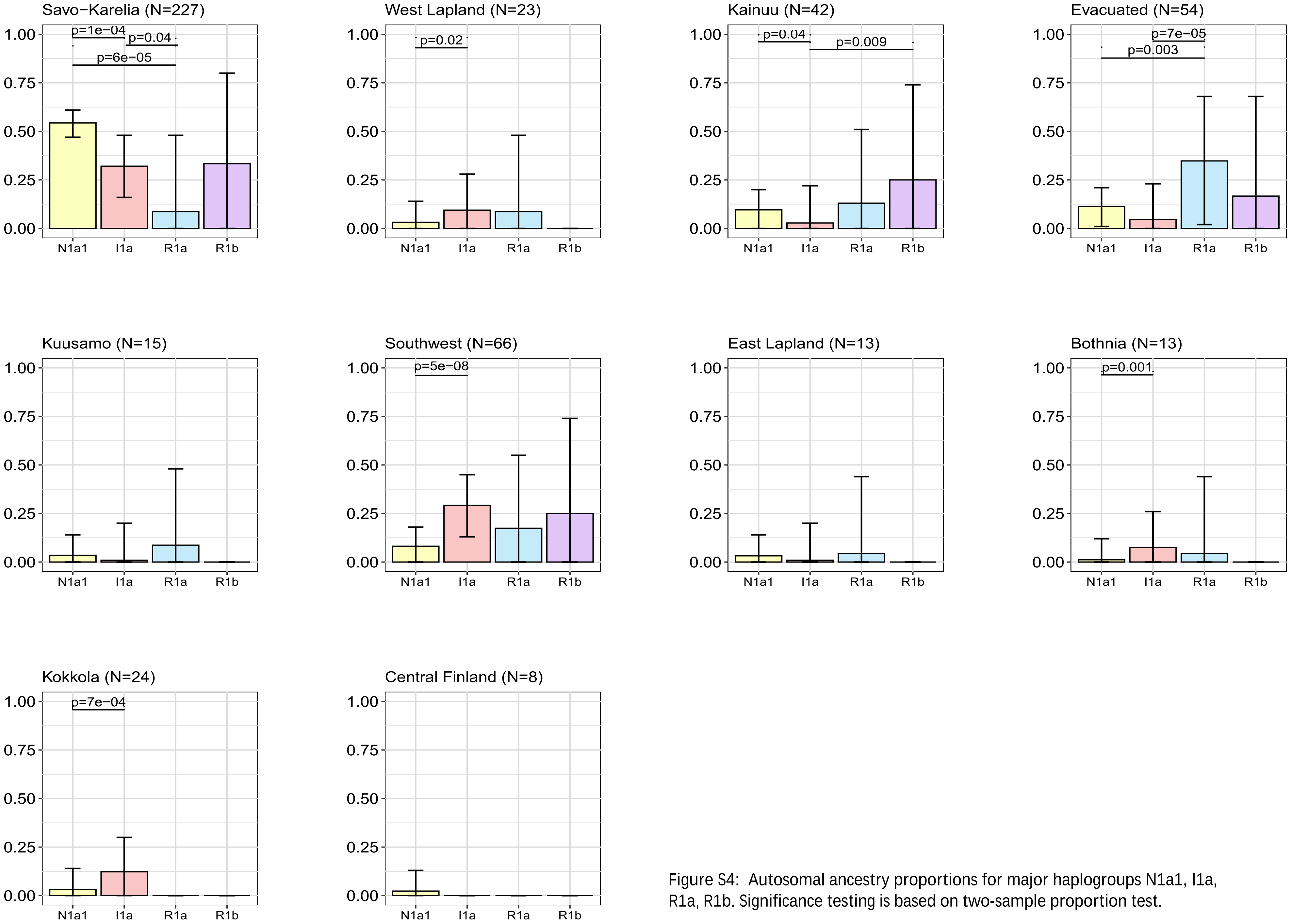


Figure S4: Autosomal ancestry proportions for major haplogroups N1a1, I1a, R1a, R1b. Significance testing is based on two-sample proportion test.

**Figure S5**

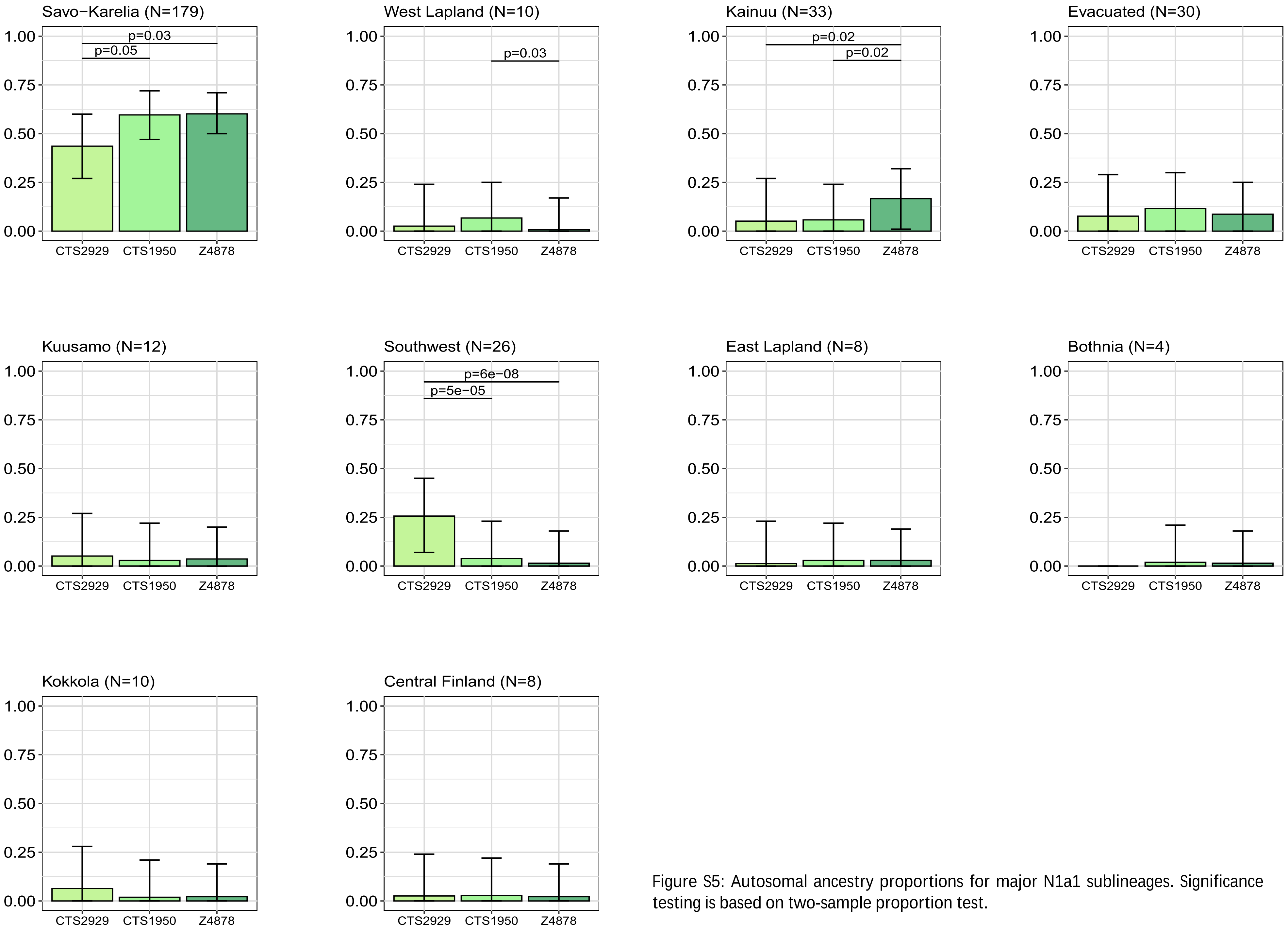


Figure S5: Autosomal ancestry proportions for major N1a1 sublineages. Significance testing is based on two-sample proportion test.
